# HIV-1 neutralization tiers are not relevant for inhibitors targeting the pre-hairpin intermediate

**DOI:** 10.1101/2022.10.06.511062

**Authors:** Benjamin N. Bell, Theodora U. J. Bruun, Natalia Friedland, Peter S. Kim

## Abstract

HIV-1 strains are categorized into one of three neutralization tiers based on the relative ease by which they are neutralized by plasma from HIV-1 infected donors not on antiretroviral therapy; tier-1 strains are particularly sensitive to neutralization while tier-2 and tier-3 strains are increasingly difficult to neutralize. Most broadly neutralizing antibodies (bnAbs) previously described target the native prefusion conformation of HIV-1 Envelope (Env), but the relevance of the tiered categories for inhibitors targeting another Env conformation, the pre-hairpin intermediate, is not well understood. Here we show that two inhibitors targeting distinct highly-conserved regions of the pre-hairpin intermediate have strikingly consistent neutralization potencies (within ∼100-fold for a given inhibitor) against strains in all three neutralization tiers of HIV-1; in contrast, best-in-class bnAbs targeting diverse Env epitopes vary by more than 10,000-fold in potency against these strains. Our results indicate that antisera-based HIV-1 neutralization tiers are not relevant for inhibitors targeting the pre-hairpin intermediate and highlight the potential for therapies and vaccine efforts targeting this conformation.

## Introduction

HIV-1 strains are routinely categorized into neutralization tiers based on the relative ease by which they are inhibited by HIV-1 antisera^1,2^, most likely by antibodies targeting Env, the only viral protein on the surface of HIV-1 virions^3^ (Fig. 1A). Tiered classification helps distinguish unusually sensitive strains in tier 1, such as those that are laboratory-adapted, from more difficult-to-neutralize strains in tiers 2 and 3 that are better representatives of currently circulating strains^1,2^ (Fig. 1A). This tiered system has also been used to systematically compare potential HIV-1 immunogens. A tier-1 neutralizing antibody response can be readily elicited by monomeric gp120 immunogens, but is not protective^4,5^; accordingly, eliciting a tier-2 neutralizing response has become the standard for evaluating prospective HIV-1 vaccine candidates^1,2,6–8^. Recent efforts to precisely categorize strains by a continuous neutralization index^2^ instead of in discrete tiers (i.e., 1A, 1B, 2, 3) illustrate the utility of this tiered system to HIV-1 research and vaccine development.

**Figure 1:**
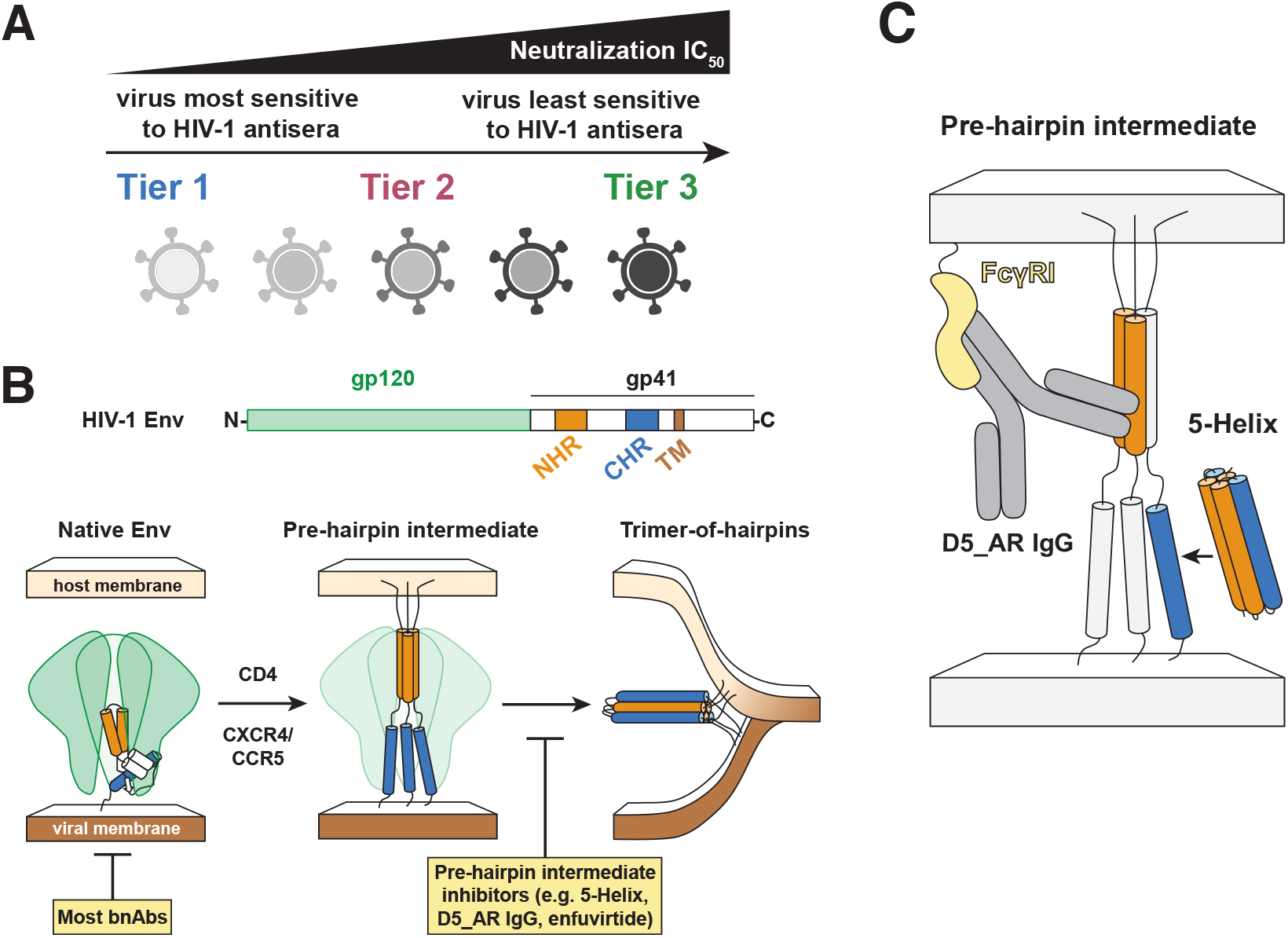
The HIV-1 pre-hairpin intermediate represents an additional neutralization target. **(A)** HIV-1 strains are organized into neutralization tiers based on their sensitivity to patient sera: tier-1 strains are particularly sensitive to neutralization, while tier-2 and tier-3 strains are more resistant. **(B)** HIV-1 Env (linear schematic), is a transmembrane (TM) viral protein composed of a trimer of gp120/gp41 heterodimers. The native, prefusion conformation of Env (left) engages cell-surface receptors CD4 and CCR5/CXCR4. The gp41 NHR and CHR are critical regions in membrane fusion exposed in the pre-hairpin intermediate (center) before collapse into a stable trimer-of-hairpins structure (right). Env-directed inhibitors target either the native prefusion conformation (left) or the pre-hairpin intermediate (center). Most bnAbs target the native prefusion state, while fusion inhibitors like 5-Helix, D5_AR IgG, and enfuvirtide prevent the transition between the pre-hairpin intermediate and the trimer-of-hairpins structure. Cartoons are based on high-resolution crystal and cryo-EM structures (PDB codes: 5FUU^55^, 6MEO^56^, 1AIK^57^, 2X7R^58^), and a common model for the pre-hairpin intermediate^10,11,32,59–61^. (**C**) The pre-hairpin intermediate inhibitors D5_AR IgG and 5-Helix target the NHR and CHR, respectively. D5_AR IgG binds a highly conserved hydrophobic pocket in the NHR trimeric coiled-coil (orange), and recent work has shown that the neutralization potency of D5_AR IgG is greatly enhanced > 1,000-fold when target cells express the cell-surface receptor FcγRI (yellow)^28,29^. 5-Helix is an engineered protein designed to mimic the post-fusion trimer-of-hairpins structure and binds the highly conserved α-helical face of the CHR (blue)^30^. Highly conserved residues are highlighted in red.

The target of antibody-mediated neutralization, HIV-1 Env, undergoes a series of conformational changes upon receptor and co-receptor binding that enable viral membrane fusion through the formation of a trimer-of-hairpins conformation^9–11^ (Fig. 1B). Most bnAbs that have been described previously target the native state of Env^12–14^, though some bnAbs are reported to bind additional Env conformations^13^. While such bnAbs have been challenging to elicit by HIV-1 vaccine candidates^6,8,15,16^, major efforts are underway to harness them for passive immunization campaigns including the recently reported Antibody Mediation Prevention trials^17–20^, which showed the bnAb VRC01 could prevent HIV-1 acquisition in healthy volunteers, but only for ∼30% of viral strains encountered in the cohort^19^. Although these trials and others show the feasibility of targeting prefusion Env in HIV-1 vaccine and therapeutic approaches, they nevertheless highlight the remaining challenges of HIV-1 prevention.

In addition to the native prefusion Env conformation, the pre-hairpin intermediate of Env can also be targeted by HIV-1 therapeutics and vaccine candidates, including by peptides^21–26^, antibodies^27–29^, and designed proteins^30^ (Fig. 1B). Binding to the gp41 N- or C-heptad repeats (NHR and CHR, respectively) in the pre-hairpin intermediate prevents formation of the trimer-of-hairpins (Fig. 1B) and is a validated inhibitory mechanism^9,10^, exemplified by the FDA-approved fusion inhibitor Fuzeon™/enfuvirtide^25,26^. Both the NHR and CHR are compelling targets, primarily due to their high sequence conservation among diverse viral clades; indeed, many positions in these regions have nearly 100% sequence identity among all known HIV-1 strains^10,11,31,32^. The NHR-targeting antibody D5_AR IgG was recently shown to weakly neutralize all strains in the tier-2 Global Reference Panel (at concentrations < 100 μg/mL)^29,33^ but with dramatic potentiation (> 1,000-fold) in the presence of FcγRI receptors^28,29^ (Fig. 1C). The CHR-targeting designed protein called 5-Helix binds highly conserved CHR residues that form an α-helical conformation^31^ (Fig. 1C); accordingly, 5-Helix has potent neutralizing activity at low nanomolar concentrations across distinct viral clades, though only a handful of viral strains have been examined to date^30,34–37^.

Despite the widespread use of the neutralization tier system to evaluate clinical approaches for HIV-1, it remains poorly understood whether such tiers predict resistance and sensitivity to inhibitors targeting the pre-hairpin intermediate^38^. Here, we determined how the neutralization potencies of D5_AR IgG and 5-Helix compare among viral strains across neutralization tiers. We ranked a panel of 18 viral strains according to neutralization tier using pooled HIV-1 antisera (HIVIG) and evaluated the potency of bnAbs targeting various Env epitopes that are being evaluated in passive immunization clinical trials: VRC01^14,19^, PGDM1400^20,39^, 3BNC117^18,40^, and 10-1074^17,41^. We find that D5_AR IgG and 5-Helix have broad and consistent neutralization potencies across neutralization tiers and viral clades (within ∼100-fold), in marked contrast to the best-in-class bnAbs targeting prefusion Env epitopes, which vary by over 10,000-fold in potency. These results suggest that HIV-1 neutralization tiers as they are currently defined are not relevant to inhibitors targeting the pre-hairpin intermediate and further highlight the appeal of HIV-1 therapeutics and vaccines that target the Env pre-hairpin intermediate.

## Results

### Inhibitors targeting the pre-hairpin intermediate have broad activity across neutralization tiers

HIV-1 neutralization tiers separate strains that are more sensitive to inhibition by sera (tier 1) from those that are more resistant to neutralization (tiers 2 and 3) but have not been evaluated in the context of inhibitors targeting the pre-hairpin intermediate. To readily compare neutralization activities across tiers and between different inhibitors and bnAbs, we first produced a panel of HIV-1 Env-pseudotyped lentiviruses representing strains that span clades and neutralization tiers including the tier-2 Global Reference Panel^33^ and select tier-1 and tier-3 strains (Table 1). We also chose to employ the widely used TZM-bl cell line, which has become the standard for assessing HIV-1 serum neutralization^42–44,33,45^.

**Table 1:** Neutralization potencies of pre-hairpin intermediate inhibitors 5-Helix and D5_AR IgG against HIV-1 panel.

| Strain | Tier | Clade | 5-Helix, IC<sub>50</sub> (µg/mL) |  | D5_AR IgG, IC<sub>50</sub> (µg/mL)<sup>29</sup> |  | D5_AR IgG, IC<sub>50</sub> (µg/mL);<br>FcγRI-expressing cells |  |
| --- | --- | --- | --- | --- | --- | --- | --- | --- |
|  |  |  | Average | St. Dev. | Average | St. Dev. | Average | St. Dev. |
| MW965.2 | 1 | C | 0.88 | 0.38 | 2.5 | 0.04 | 0.004 | 0 |
| SF162 | 1 | B | 2.7 | 0.21 | 40 | 15 | 0.01 | 0 |
| HXB2 | 1 | B | 1.1 | 0.25 | 5.4 | 1.0 | 0.007 | 0 |
| BaL | 1 | B | 3.5 | 0.5 | 26 | 8.3 | 0.045 | 0.0071 |
| SS1196.1 | 1 | B | 1.9 | 1.4 | 23 | 10 | 0.025 | 0.0071 |
| 25710 | 2 | C | 1.9 | 0.24 | 4.6 | 0.7 | 0.014 | 0 |
| TRO.11 | 2 | B | 1.5 | 0.25 | 54 | 14 | 0.025 | 0.0071 |
| BJOX2000 | 2 | CRF07_BC | 1.4 | 0.22 | 46 | 6.0 | 0.008 | 0.0014 |
| X1632 | 2 | G | 3.0 | 0.72 | 66 | 8.0 | 0.035 | 0.0071 |
| CE1176 | 2 | C | 2.8 | 0.71 | 79 | 10 | 0.15 | 0.071 |
| 246F3 | 2 | AC | 2.3 | 0.35 | 28 | 1.9 | 0.015 | 0.0071 |
| CH119 | 2 | CRF07_BC | 2.2 | 0.29 | 37 | 4.7 | 0.06 | 0.028 |
| CE0217 | 2 | C | 2.2 | 0.069 | 27 | 6.0 | 0.025 | 0.0071 |
| CNE55 | 2 | CRF01_AE | 5.2 | 0.87 | 26 | 2.2 | 0.02 | 0.014 |
| TRJO | 3 | B | 1.6 | 0.38 | 29 | 14 | 0.065 | 0.0071 |
| 33-7 | 3 | CRF02_AG | 1.2 | 0.18 | 29 | 0.21 | 0.02 | 0 |
| PVO.4 | 3 | B | 5.9 | 0.52 | 34 | 0.27 | 0.035 | 0.0071 |
| 253-11 | 3 | CRF02_AG | 0.51 | 0.099 | 12 | 1.0 | 0.0065 | 0.00071 |

Although 5-Helix neutralization activity has been reported using a number of cell-cell fusion^30,34^ and virus-cell fusion^34–37^ assays, no reports to date have employed TZM-bl cells. Therefore, we first sought to determine 5-Helix potencies using TZM-bl cells and corroborate our findings using HOS-CD4-CCR5 target cells for which 5-Helix potencies have been published^34–37^. We validated our 5-Helix preparations biochemically (SI Fig. 1A-G) and found good agreement between our half-maximal inhibitory concentration (IC_50_) value using HOS-CD4-CCR5 cells and HXB2 Env-pseudotyped lentivirus (8.2 ± 4.1 nM; SI Fig. 2A), compared to a previously published value using this strain and cell line (11 ± 1.2 nM^34^). We observed a reduction in 5-Helix neutralization potency using TZM-bl cells compared to HOS-CD4-CCR5 cells across multiple strains (SI Fig. 2A), in line with similarly enhanced activity in HOS-CD4-CCR5 over TZM-bl target cells observed with NHR-targeting inhibitors HK20, D5, and Fuzeon™/enfuvirtide^27^. Such discrepancies across neutralization assay format are well documented^46^, likely due to differences in viral membrane fusion in these cell lines.

**Figure 2:**
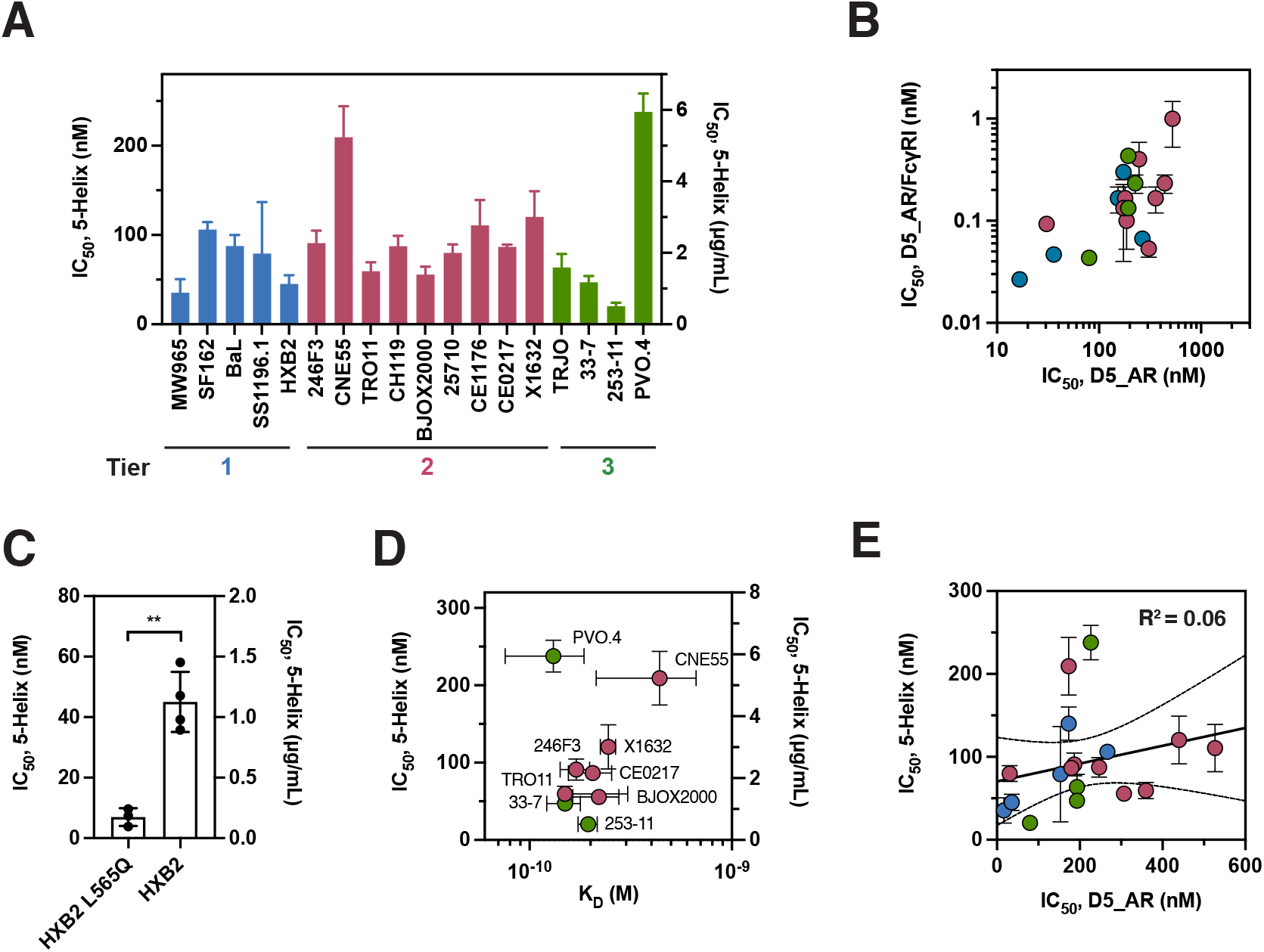
Inhibitors targeting the pre-hairpin intermediate have broad activity across neutralization tiers. (**A**) CHR-targeting inhibitor 5-Helix shows activity across a roughly ten-fold range against viral strains in neutralization tiers 1/2/3. In panels A-E, IC_50_ values from TZM-bl neutralization assays performed in biological triplicate are plotted as mean ± standard deviation and colored by neutralization tier: tier 1 (blue), tier 2 (red), and tier 3 (green). (**B**) NHR-targeting D5_AR IgG shows activity against viral strains in neutralization tiers 1/2/3 that is potentiated roughly 1,000-fold for cells expressing FcγRI. Mean IC_50_ values ± standard deviation of repeated neutralization assays done in at least biological triplicate using either TZM-bl (x-axis) or TZM-bl/FcγRI (y-axis) cells are shown. (**C**) HIV-1 strains with identical CHR sequences (HXB2 and HXB2 L565Q) show differences in 5-Helix sensitivity. Two-tailed unpaired t-tests were performed with the indicated p-value (**p < 0.01). (**D**) Increased binding of 5-Helix to CHR sequences measured by biolayer interferometry does not correspond to increased sensitivity to inhibition by 5-Helix. Reported K_D_ values (mean ± standard deviation) were determined from repeated experiments (n=2) for each peptide across a range of concentrations of 5-Helix. (**E**) Sensitivity to 5-Helix does not predict sensitivity to D5_AR IgG. IC_50_ values of 5-Helix and D5_AR IgG fall over notably different ranges and show only a modest association. A linear regression (black) with 95% confidence bands (dashed) has a correlation coefficient of R^2^ = 0.05.

We next evaluated 5-Helix neutralization potency against strains across HIV-1 neutralization tiers. 5-Helix had broad activity across neutralization tiers and viral clades in a narrow range of IC_50_ values (∼10-fold), including potent inhibition of three of the four tier-3 strains tested (< 80 nM or < 2 μg/mL; Fig. 2A, Table 1). Indeed, one tier-3 strain, 253-11, characterized as being unusually resistant to neutralization^47^, was especially sensitive to inhibition by 5-Helix (Fig. 2A, Table 1). To validate that the observed sensitivity to 5-Helix was not an artifact of our tier-3 pseudovirus preparations, we tested these strains against the bnAb 10E8v4 IgG1 and found close agreement with published values^48^ (SI Fig. 2B), indicating that this sensitivity to 5-Helix is genuine.

As the pre-hairpin intermediate can be inhibited by targeting either the CHR or the NHR, we next used our pseudovirus panel to evaluate the neutralization potencies of the NHR-targeting antibody D5_AR IgG, which was first described to weakly neutralize all strains in the tier-2 Global Reference Panel^29,33^. We found similar potencies for D5_AR IgG and also saw at least 1,000-fold increases in neutralization potency in the presence of the cell-surface receptor FcγRI as previously described^28,29^ (Fig. 2B). Like 5-Helix, D5_AR IgG also showed a narrow range of IC_50_ values (∼100-fold) that did not correlate well with neutralization tier (Fig. 2B).

As neutralization tier does not seem to account for variable sensitivity to 5-Helix and D5_AR IgG, we considered whether differing binding affinities of 5-Helix to CHR sequences correspond to differing sensitivities among strains. The NHR mutation L565Q does not affect the CHR sequence; thus, one might expect 5-Helix would show similar neutralization potency in the presence or absence of the mutation. However, the L565Q virus showed increased sensitivity to 5-Helix (Fig. 2C), in line with previous reports^37,49^, indicating that epitope sequence is not the only determinant of sensitivity to 5-Helix. To further probe how sequence diversity impacts 5-Helix efficacy, we next assayed 5-Helix binding to nine peptides representing the CHR sequences of some of the strains tested across viral clades in our panel (Fig. 2D, Supplementary Table 1). Due to the extremely high affinity of 5-Helix for the CHR (K_D_ = 0.6 pM in the case of HXB2^34^), we measured binding affinity by biolayer interferometry using steady-state values after 1 h of association in the presence of 1 M guanidine hydrochloride (SI Fig. 3A-I). Consistent with our observations of HXB2 vs. HXB2 L565Q (Fig. 2C), there was not a strong correlation between 5-Helix binding affinity for various CHR sequences and IC_50_ values (Fig. 2D).

**Figure 3:**
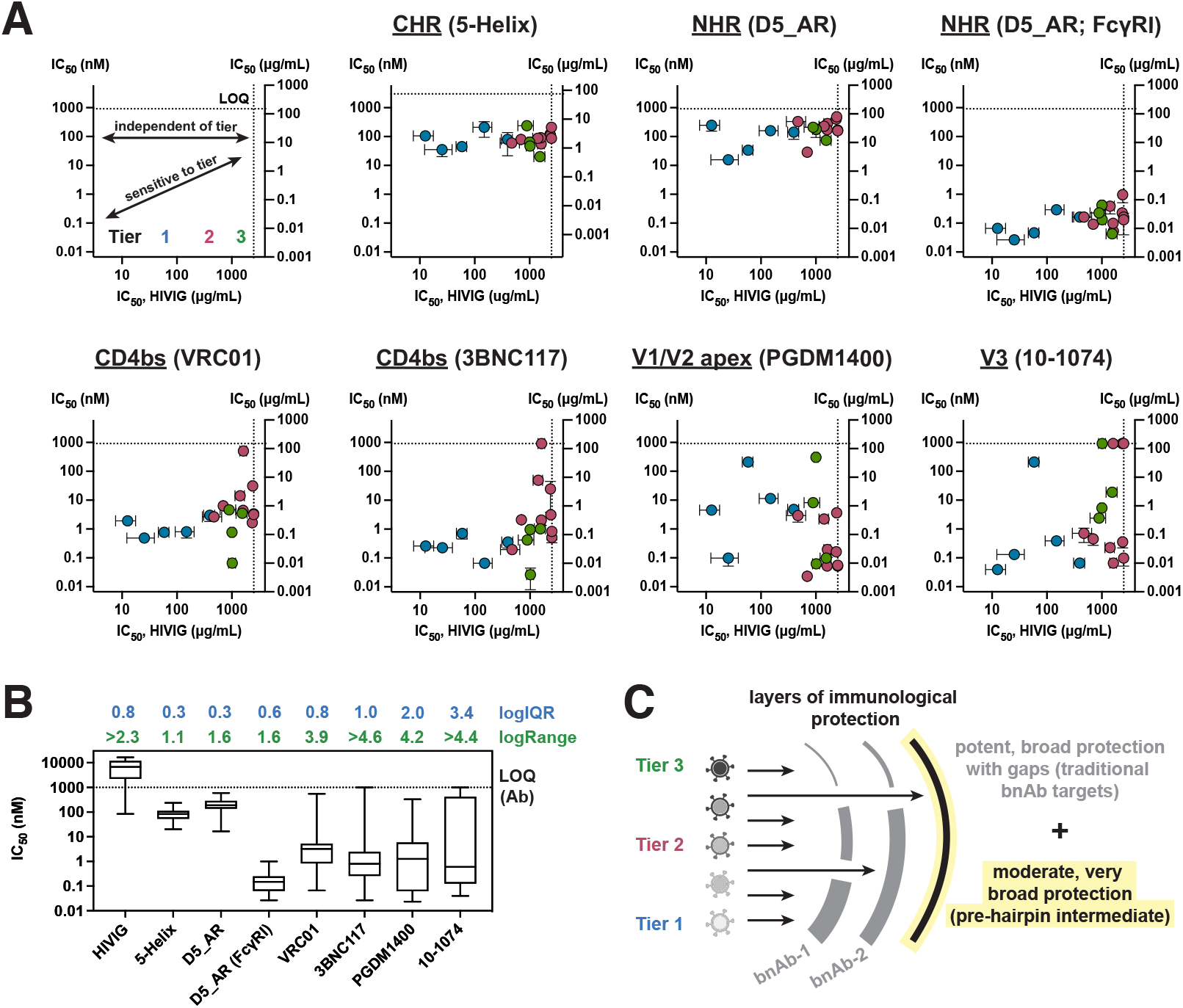
Pre-hairpin intermediate inhibitors show highly consistent and broad HIV-1 neutralization independent of tier, unlike best-in-class bnAbs. (**A**) 5-Helix and D5_AR IgG show highly consistent neutralization activity across tiers. Neutralization IC_50_ values were determined against a panel of tier-1/2/3 strains for immunoglobin purified from pooled HIV-1 sera (HIVIG) and four bnAbs employed in clinical trials VRC01, 3BNC117, PGDM1400, and 10-1074. Using HIVIG as an indicator of tier, only 5-Helix and D5_AR IgG (in TZM-bl cells ± FcγRI expression) show highly consistent activity among strains across tiers. The limit-of-quantitation (LOQ) is indicated by dashed lines. Values are plotted as mean ± standard deviation and colored by neutralization tier: tier 1 (blue), tier 2 (red), and tier 3 (green). Epitopes of each inhibitor are indicated: CD4 binding site (CD4bs), gp120 V1/V2 apex, and gp120 V3 loop. (**B**) Box-and-whisker plots of IC_50_ values in panel A show 5-Helix and D5_AR IgG have narrower ranges than HIVIG or best-in-class bnAbs. The logarithms of the interquartile range (logIQR) and the range (logRange) are reported. The limit-of-quantitation (LOQ) for the antibody neutralization datasets is indicated by the dashed line. (**C**) Proposed model for incorporating inhibitors targeting the pre-hairpin intermediate into current HIV-1 prevention approaches. Viruses from different neutralization tiers show different susceptibility to current best-in-class bnAbs, but similar vulnerability to inhibitors targeting the pre-hairpin intermediate. High neutralization potency is represented as thicker layers of protection with no potency represented as a gap in the layer. Eliciting antibodies that neutralize via the pre-hairpin intermediate may provide an additional layer of protection that would be useful even if moderate but very broad.

As CHR sequence alone does not fully determine strain sensitivity to 5-Helix, we next considered whether there is a correlation between strains that were sensitive to 5-Helix and those sensitive to D5_AR IgG. Previous studies using 5-Helix and CHR-peptide mutants proposed that the CHR and NHR have distinct windows of vulnerability and accessibility in the pre-hairpin intermediate^34^. Indeed, we found poor agreement between strains that were sensitive to inhibition by 5-Helix and those sensitive to inhibition by D5_AR IgG, with poor linear correlation (R^2^ = 0.06; Fig. 2E). Additionally, D5_AR IgG IC_50_ values span a slightly larger range than those of 5-Helix (100-fold versus 10-fold; Fig. 2E, Table 1). Therefore, strain sensitivity to pre-hairpin intermediate inhibitors is not explained by differences in target binding affinity (Fig. 2C-D), nor do the NHR and CHR have equal susceptibility (Fig. 2E).

### Pre-hairpin intermediate inhibitors show highly consistent and broad HIV-1 neutralization independent of tier, unlike best-in-class bnAbs

We sought to determine whether the broad, narrow range of potencies we observed for 5-Helix and D5_AR IgG were unique; accordingly, we assayed our 18-virus panel in neutralization assays using best-in-class bnAbs currently employed in clinical trials: VRC01^19^ and 3BNC117^18^ (targeting the CD4 binding site in gp120), PGDM1400^20^ (targeting the gp120 V1/V2 apex), and 10-1074^17^ (targeting the gp120 V3 loop). We used HIVIG (purified immunoglobin from pooled HIV-1 patient sera) as a quantitative representation of neutralization tier, as similar pools of patient sera are used to delineate these tiers (Table 2)^2,38^.

**Table 2:**
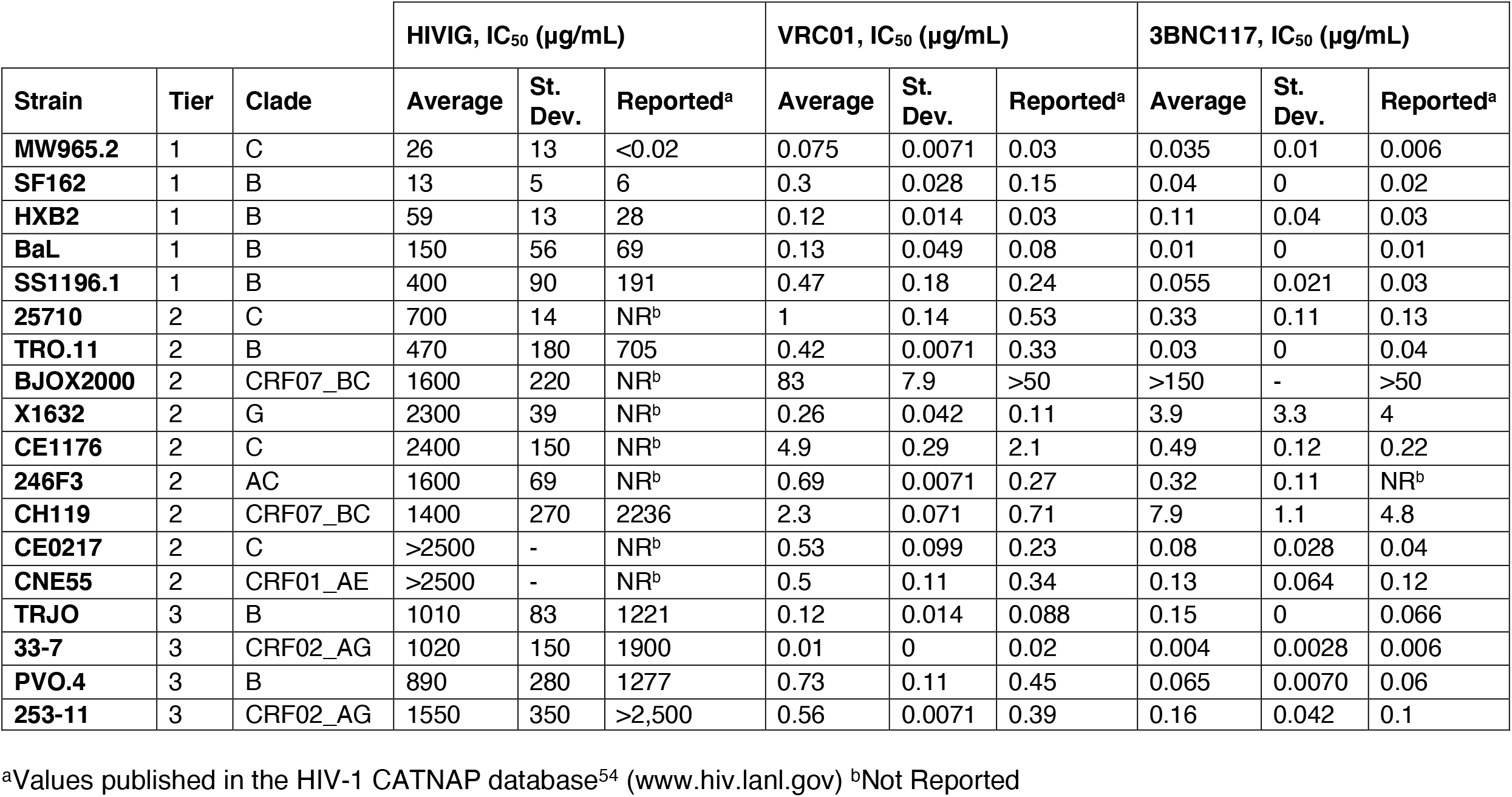
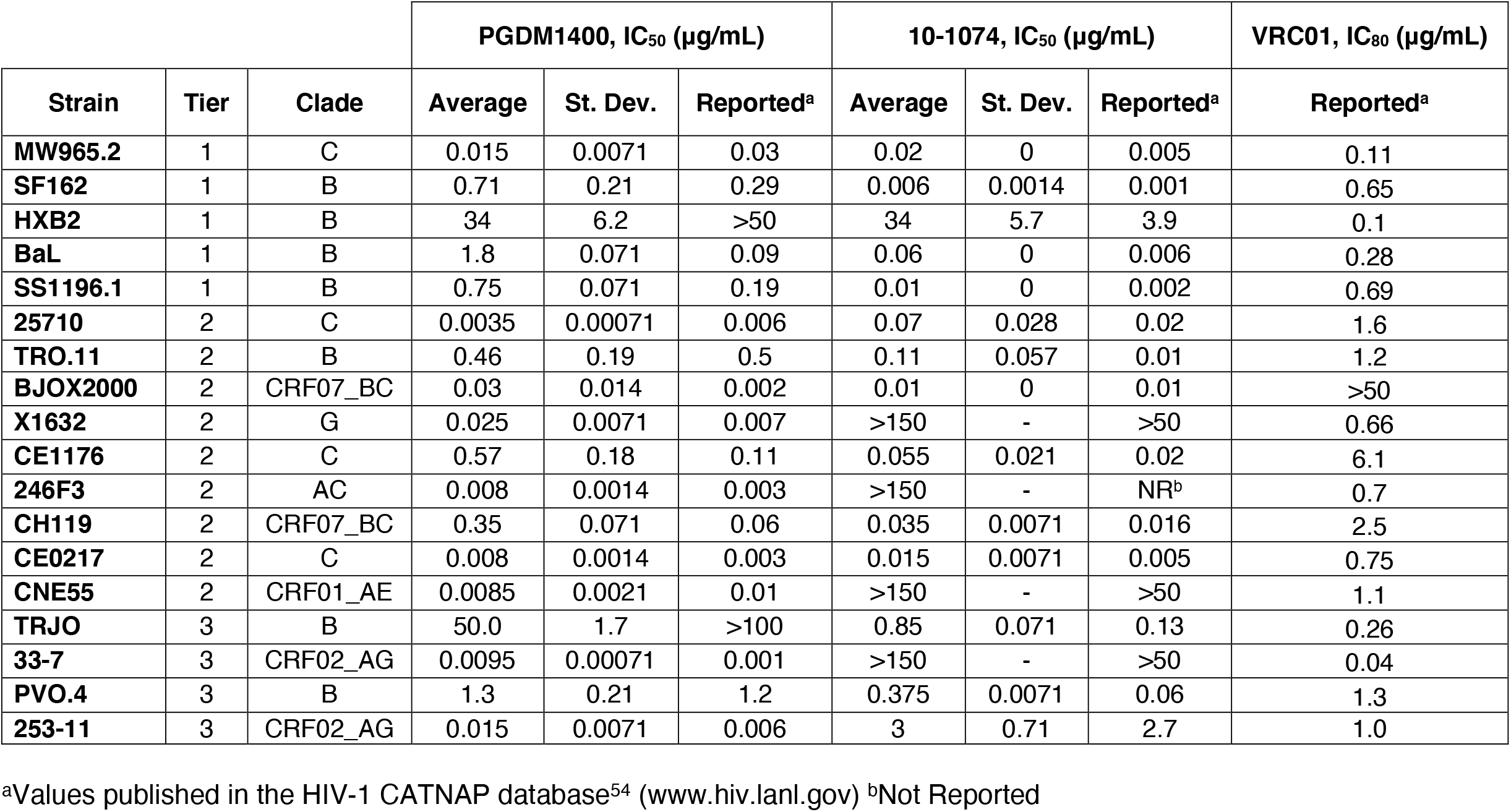
Neutralization potencies of best-in-class bnAbs against HIV-1 panel.

We found striking differences between the bnAbs evaluated here and 5-Helix or D5_AR IgG (Fig. 3A). The neutralization potencies of 5-Helix and D5_AR IgG did not vary substantially across a wide range of HIVIG potencies (13 to > 2500 μg/mL), whereas the four bnAbs varied widely (Fig. 3A). We quantified the variation in IC_50_ values by examining the log-values of the interquartile range (IQR) as well as the total range of the data (Fig. 3B). While 5-Helix and D5_AR IgG IC_50_ values varied by up to two logs (∼100-fold), the clinical bnAbs tested showed ranges of ∼10,000-fold (VRC01) to nearly 100,000-fold (3BNC117, 10-1074; Fig. 3A,B). Taken together, these results show that neutralization tiers have minimal predictive power for inhibition by 5-Helix and D5_AR IgG; further, such inhibitors show extremely narrow ranges of IC_50_ values unlike best-in-class bnAbs currently employed in clinical studies.

## Discussion

HIV-1 neutralization tiers are a powerful tool to evaluate novel strains and vaccine candidates^33,38^ as the neutralization activity of bnAbs and vaccine sera can vary substantially against tier-2 and tier-3 strains. Here we show that 5-Helix and D5_AR IgG, two inhibitors targeting highly conserved gp41 epitopes in the pre-hairpin intermediate, can have broad inhibitory activity against strains in all three neutralization tiers of HIV-1, including particularly resistant tier-3 strains (Fig. 2A, Table 1). Although 5-Helix and D5_AR IgG neutralization potencies are modest (10-1,000 nM, Fig. 3A), they show very narrow variation across tiers (less than ∼100-fold), unlike best-in-class bnAbs currently employed in clinical trials (more than 10,000-fold, Fig. 3A,B)^17–20^. Notably, even though D5_AR is a generally weak inhibitor of HIV-1 infection, dramatic potentiation observed in the presence of FcγRI suggests more robust neutralization activity might occur *in vivo*^29,50^. Further, D5_AR IgG showed superior neutralization potency compared to the bnAbs studied here to some tier-2 (BJOX2000, X1632, 246F3, CNE55) and tier-3 (TRJO, 33-7) strains (Fig. 3A, Table 1, Supplementary Table 1). Overall, these results indicate that HIV-1 neutralization tiers as they are currently defined are not relevant for inhibitors targeting the pre-hairpin intermediate.

If neutralization tiers do not predict sensitivity to pre-hairpin intermediate inhibitors, what factors may be at play? Differences in the binding affinities between 5-Helix and distinct CHR sequences did not account for differences in sensitivity to inhibition by 5-Helix (Fig. 2C,D). Targeting the NHR and CHR do not appear to be equivalent: we observed a wider range of IC_50_ values for D5_AR IgG compared to 5-Helix, and strains more resistant to 5-Helix were not necessarily more resistant to D5_AR IgG (Fig. 2E). The pre-hairpin intermediate is a transient structure, and its lifetime may vary among strains; it is therefore possible that different fusion kinetics could affect inhibitor access and efficacy. Indeed, mutations in the gp41 NHR that showed resistance to Fuzeon™/enfuvirtide conferred delayed viral membrane fusion kinetics^49,51^, and such mutations sensitize strains to inhibition by 5-Helix^37^ (e.g., HXB2 L565Q, Fig. 2C). Further, differences in steric^52,53^ and kinetic^34–36^ accessibility of the NHR and CHR has been documented using 5-Helix and CHR peptide variants. These hypotheses warrant future study to identify the determinants of differing sensitivity to pre-hairpin intermediate inhibitors.

Current bnAbs used in the clinic have shown the promise of passive immunization approaches^17–20^, but the large fraction of resistant virus encountered already in these trials highlights the challenge of gaps in which these bnAbs do not provide protection (Fig. 3C). Indeed, in recent trials using VRC01, nearly 70% of participants were infected by strains with at least some resistance to VRC01 neutralization (defined as IC_80_ ≥ 1 μg/mL)^19^. Pharmacokinetic estimates suggest participants had VRC01 serum concentrations of at least 10 μg/mL throughout the study period, setting a potential threshold of protection around ten times the IC_80_ value for a given strain. Based on these considerations, 8 of the 13 tier-2/3 strains in our panel would be predicted to infect VRC01-trial participants^19,54^ (Table 2; last column). While current inhibitors against the pre-hairpin intermediate would not fare better than clinical bnAbs at these concentrations, the narrow range of IC_50_ values we observe suggest these targets have the potential to provide a very broad layer of protection not afforded by current bnAbs (Fig. 3C).

Developing a vaccine that protects against clinical HIV-1 isolates (i.e., tier-2 and tier-3 strains) remains a significant challenge. Our results demonstrate that inhibitors targeting the pre-hairpin intermediate conformation of HIV-1 Env are strikingly less influenced by neutralization tier compared to best-in-class bnAbs. This suggests that vaccine strategies that successfully elicit antibodies targeting the pre-hairpin intermediate could provide an especially broad layer of protection. Further, this protection could be even more effective if FcγRI enhancement proves to be relevant *in vivo*. Overall, this work highlights the appeal of creating an HIV-1 vaccine that can elicit anti-CHR and/or anti-NHR neutralizing antibodies mimicking the broad binding and neutralization properties of 5-Helix and D5_AR IgG.

**Supplementary Figure 1:**
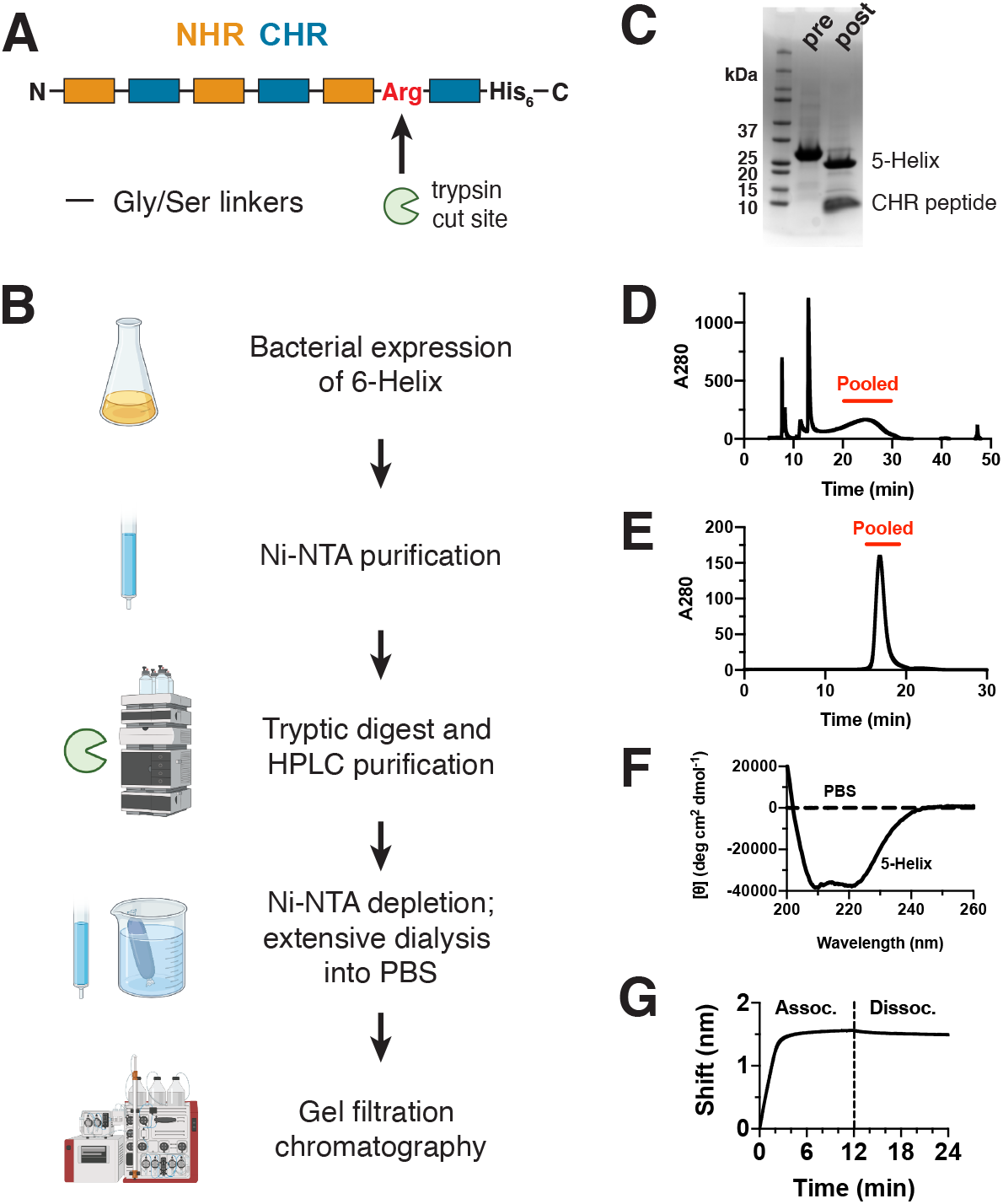
Validation of 5-Helix purification scheme. (**A**) 5-Helix is generated from a 6-Helix construct in which three NHR and three CHR sequences are connected via Gly/Ser linkers with a C-terminal His_6_ purification tag. An Arg residue in the final linker is sensitive to trypsin digestion to produce 5-Helix and a CHR peptide. (**B**) Overview of 5-Helix purification scheme: 6-Helix is first expressed in bacteria and purified by Ni-NTA affinity chromatography. Following limited tryptic digest, reverse phase chromatography is used to isolate 5-Helix. Fractions are diluted with urea and residual 6-Helix and CHR peptide are removed by binding to Ni-NTA resin. After extensive dialysis into PBS, 5-Helix is purified by gel filtration chromatography. (**C**) SDS-PAGE analysis of 6-Helix protein pre- and post-tryptic digest. (**D**) Representative HPLC trace of trypsin-digested 6-Helix with pooled fractions indicated. (**E**) Representative gel filtration trace of refolded 5-Helix with pooled fractions indicated shows monodisperse population of 5-Helix. (**F**) Representative circular dichroism shows purified 5-Helix is folded with high α-helicity. (**G**) Representative biolayer interferometry curve with association and dissociation steps indicated shows 50 nM 5-Helix binds to CHR peptide HXB2.

**Supplementary Figure 2:**
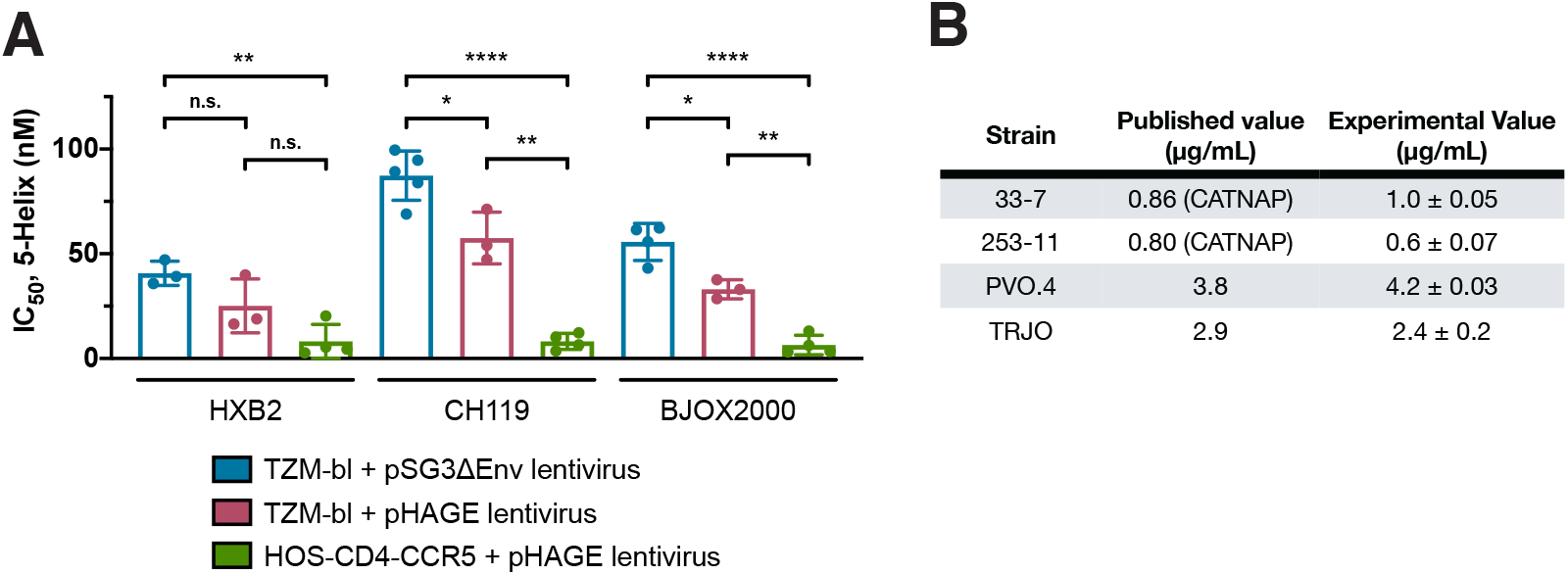
Inhibition by 5-Helix in TZM-bl assay agrees well with published observations. (**A**) IC_50_ values of 5-Helix activity differ from three assay formats against three HIV-1 strains: HXB2 (tier-1B, clade B), CH119 (tier-2, CRF07_BC), and BJOX2000 (tier-2, CRF07_BC). Mean ± standard deviation of repeated experiments done in at least biological triplicate are shown. Two-tailed unpaired t-tests were performed with indicated p-values (n.s. = not significant; *p < 0.05; **p<0.01; ***p<0.001; ****p<0.0001). (**B**) Inhibition of tier-3 viral strains by 10E8v4 IgG1 show good agreement with published values^48^ (including CATNAP database). Mean ± standard deviation of repeated experiments done in biological duplicate are shown.

**Supplementary Figure 3:**
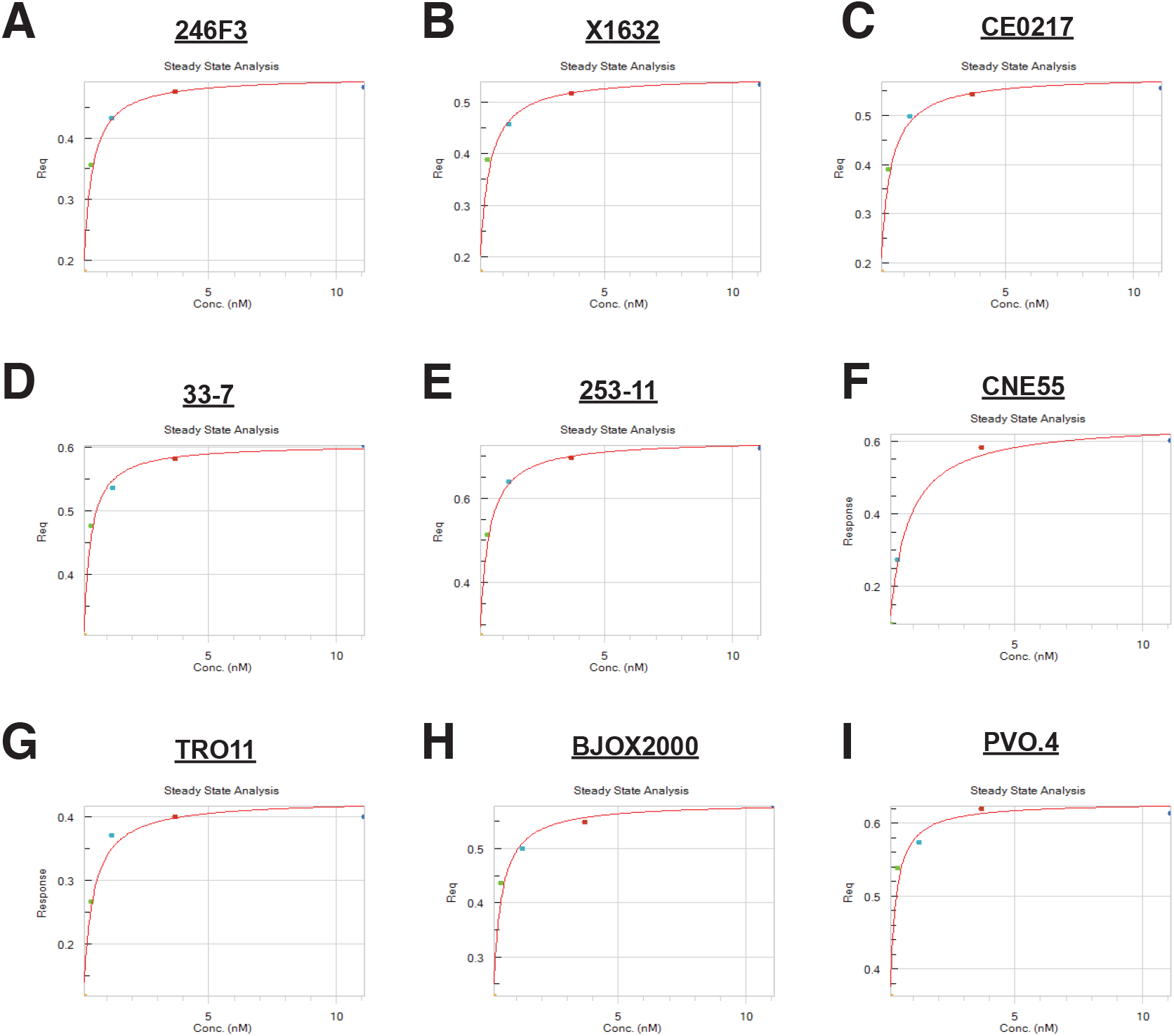
Representative binding steady state binding values for 5-Helix to CHR peptides. (**A-I**) Steady state values after a one-hour association step in biolayer interferometry experiments used to calculate K_D_ values of 5-Helix binding to CHR peptides from the indicated viral strain.

**Supplementary Table 1:**
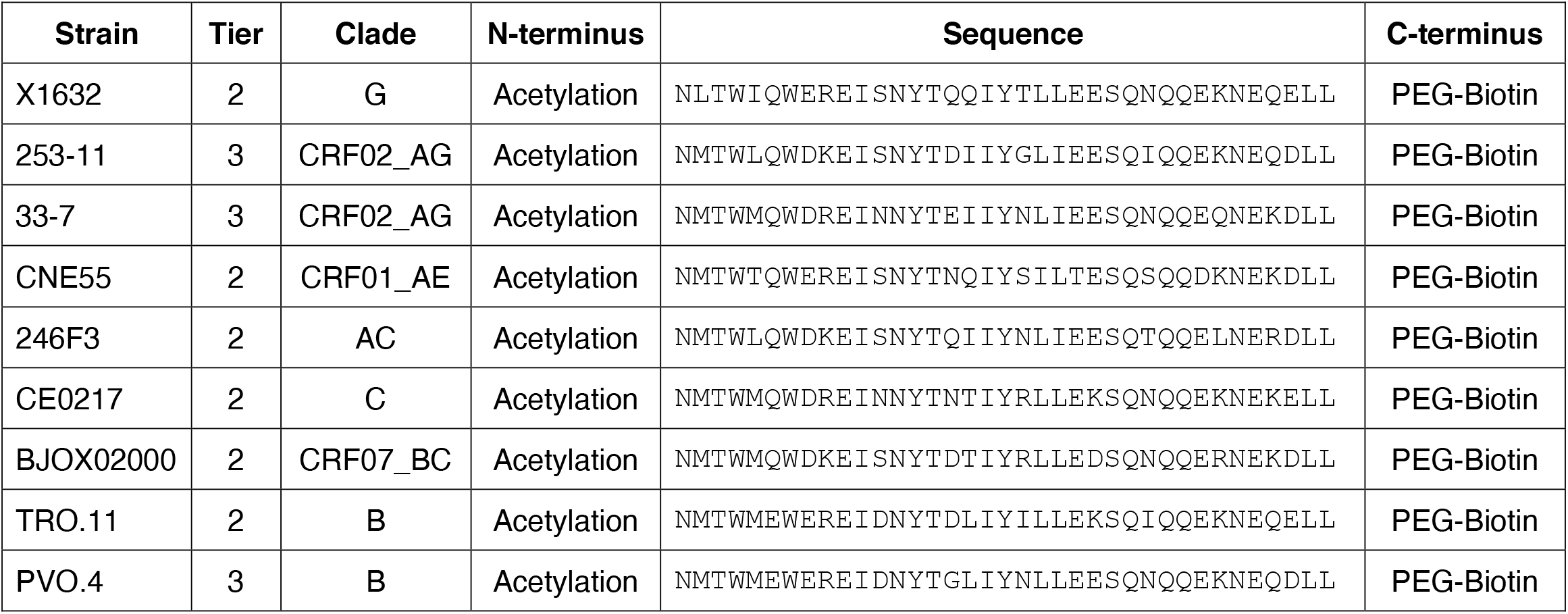
CHR peptides used in binding experiments.

## Materials and Methods

### NIH HIV Reagents

The following reagents were obtained through the NIH HIV Reagent Program, Division of AIDS, NIAID, NIH: (1) Polyclonal Anti-Human Immunodeficiency Virus Immune Globulin, Pooled Inactivated Human Sera, ARP-3957, contributed by NABI and National Heart Lung and Blood Institute (Dr. Luiz Barbosa); (2) HOS CD4+ CCR5+ Cells, ARP-3318, contributed by Dr. Nathaniel Landau, Aaron Diamond AIDS Research Center, The Rockefeller University. (3) TZM-bl Cells, ARP-8129, contributed by Dr. John C. Kappes, Dr. Xiaoyun Wu and Tranzyme Inc.; (4) Panel of Global Human Immunodeficiency Virus 1 (HIV-1) Env Clones, ARP-12670, from Dr. David Montefiori; (5) Vector pSV7d Expressing Human Immunodeficiency Virus Type 1 (HIV-1) HXB2 Env (pHXB2-env), ARP-1069, contributed by Dr. Kathleen Page and Dr. Dan Littman; 6) Human Immunodeficiency Virus 1 93MW965.26 gp160 Expression Vector (pSVIII-93MW965.26), ARP-3094, contributed by Dr. Beatrice Hahn; (7) Human Immunodeficiency Virus 1 (HIV-1) BaL.26 Env Expression Vector, ARP-11446, contributed by Dr. John Mascola; (8) Human Immunodeficiency Virus 1 (HIV-1) SF162 gp160 Expression Vector, ARP-10463, contributed by Dr. Leonidas Stamatatos and Dr. Cecilia Cheng-Mayer; (9) Plasmid pcDNA3.1 D/V5-His TOPO© Expressing Human Immunodeficiency Virus Type 1 Env/Rev, ARP-11020, contributed by Dr. David Montefiori and Dr. Feng Gao; (10) Human Immunodeficiency Virus 1 Panel of HIV-1 Subtype A/G Env Clones, ARP-11673, contributed by Drs. D. Ellenberger, B. Li, M. Callahan, and S. Butera; (11) Panel of Human Immunodeficiency Virus Type 1 Subtype B Env Clones, ARP-11227, contributed by Drs. D. Montefiori, F. Gao, M. Li, B.H. Hahn, X. Wei, G.M. Shaw, J.F. Salazar-Gonzalez, D.L. Kothe, J.C. Kappes and X. Wu. (12) The following reagent was obtained through the NIH HIV Reagent Program, Division of AIDS, NIAID, NIH: Human Immunodeficiency Virus Type 1 (HIV-1) SG3ΔEnv Non-infectious Molecular Clone, ARP-11051, contributed by Dr. John C. Kappes and Dr. Xiaoyun Wu; (13) Vector CMV/R Containing the Human IgG1 Heavy Chain Gene for Expression of Anti-Human Immunodeficiency Virus 1 (HIV-1) gp41 Monoclonal Antibody 10E8v4 in 293-6E Cells, APR-12866, contributed by Dr. Peter Kwong; (14) Vector CMV/R Containing the Human IgG1 Light Chain Gene for Expression of Anti-Human Immunodeficiency Virus 1 (HIV-1) gp41 Monoclonal Antibody 10E8v4 in 293-6E Cells, APR-12867, contributed by Dr. Peter Kwong; (15) Anti-Human Immunodeficiency virus (HIV)-1 gp120 Monoclonal Antibody (3BNC117), ARP-12474, contributed by Dr. Michel Nussenzweig; (16) Anti-Human Immunodeficiency Virus (HIV)-1 gp120 Monoclonal antibody (10-1074), ARP-12477, contributed by Dr. Michel Nussenzweig; (17) Human Immunodeficiency Virus 1 (HIV-1) VRC01 Monoclonal Antibody Heavy Chain Expression Vector, ARP-12035, contributed by John Mascola; (18) Human immunodeficiency Virus 1 (HIV-1) VRC01 Monoclonal Antibody Light Chain Expression Vector, ARP-12036, contributed by John Mascola.

### 5-Helix expression/purification

5-Helix was recombinantly expressed and purified as previously described^30^. Briefly, a 6-Helix construct was expressed recombinantly in BL21 (DE3) *Escherichia coli* (New England Biolabs). This construct was composed of three NHR and three CHR peptides with intervening glycine/serine linkers and a C-terminal hexahistidine purification tag. The final glycine/serine linker contained an arginine residue sensitive to trypsin cleavage. *E. coli* cultures were induced at OD_600_ ∼0.6-0.8 with 1 mM IPTG and harvested after 3 h expression at 37 °C shaking at 225 rpm.

Cell pellets were lysed via sonication in Tris-buffered saline (TBS: 25 mM Tris-HCl [pH 8.0], 100 mM NaCl) and bound to 1 mL Ni-NTA agarose (Thermo Fisher Scientific) for 2 h at 4 °C with agitation. 6-Helix was subsequently eluted from the Ni-NTA resin with TBS + 250 mM imidazole [pH 8.0] following a wash with TBS + 25 mM imidazole [pH 8.0]. Eluted protein was digested with trypsin (1:200 w/w) for 15-20 min in a shaking-platform incubator at 37 °C shaking at 100 rpm.

Trypsin-digested 6-Helix protein was then purified by high-pressure liquid chromatography (HPLC) on a C18 semi-preparative column (Phenomenex) over a 38-45% acetonitrile gradient in the presence of 0.1% trifluoroacetic acid. 5-Helix-containing HPLC fractions were analyzed by SDS-PAGE and pooled. Pooled fractions were diluted with TBS and 8 M urea [pH 8.0] to a final protein concentration of ∼0.1-0.2 mg/mL and residual 6-Helix and CHR peptide were removed by binding to Ni-NTA resin for 1 h. The flow-through from this step was dialyzed overnight into PBS [pH 7.4]. Following two additional 2 h dialysis steps into PBS, 5-Helix was concentrated to 2 mg/mL and flash frozen with LN_2_ with 10% glycerol. A final gel-filtration chromatography purification step was performed using a Superdex 200 Increase 10/300 GL column (Cytiva) on a Cytiva ÄKTA Pure system immediately before use.

The 6-Helix protein sequence used to generate 5-Helix protein is: MQLLSGIVQQQNNLLRAIEAQQHLLQLTVWGIKQLQARILAGGSGGHTTWMEWDREINNYTS IHSLIEESQNQQEKNEQELLEGSSGGQLLSGIVQQQNNLLRAIEAQQHLLQLTVWGIKQLQA RILAGGSGGHTTWMEWDREINNYTSLIHSLIEESQNQQEKNEQELLEGSSGGQLLSGIVQQQNNLLRAIEAQQHLLQLTVWGIKQLQARILAGGRGGGHTTWMEWDREINNYTSLIHSLIEESQNQQEKNEQELLEGGHHHHHH. The 5-Helix protein sequence is: MQLLSGIVQQQNNLLRAIEAQQHLLQLTVWGIKQLQARILAGGSGGHTTWMEWDREINNYTSLIHSLIEESQNQQEKNEQELLEGSSGGQLLSGIVQQQNNLLRAIEAQQHLLQLTVWGIKQLQARILAGGSGGHTTWMEWDREINNYTSLIHSLIEESQNQQEKNEQELLEGSSGGQLLSGIVQQQNNLLRAIEAQQHLLQLTVWGIKQLQARILAGGR.

### Antibody expression/purification

D5_AR, 10E8v4, VRC01, PGDM1400, and 10-1074 IgG1s were expressed and purified from Expi293F cells. Expression vectors for D5_AR were generated previously^29^, expression vectors for VRC01 and 10E8v4 were sourced from the NIH HIV Reagent Program (see “NIH HIV Reagents”), and expression vectors for 10-1074 were gifted from Dr. Christopher Barnes^17,41^. PGDM1400 heavy and light chain sequences were synthesized (Integrated DNA Technologies) and cloned into a mammalian expression vector under a CMV promoter using InFusion (Takara) and sequence verified.

Expi293F cells were cultured in 33% Expi293 Expression / 66% FreeStyle Expression medium (Thermo Fisher Scientific) and grown in baffled polycarbonate shaking flasks (Triforest) at 37 °C and 8% CO_2_. Cells were grown to a density of ∼3 × 10^6^/mL and transiently transfected using FectoPro transfection reagent (Polyplus). For transfections, 0.5 μg total DNA (1:1 heavy chain to light chain plasmids) was added per mL final transfection volume to culture medium (1/10 volume of final transfection) followed by FectoPro at a concentration of 1.3 μL per mL final transfection volume and incubated at room temperature for 10 min. Transfection mixtures were added to cells, which were then supplemented with D-glucose (4 g/L final concentration) and 2-propylpentanoic (valproic) acid (3 mM final concentration). Cells were harvested 3-5 days after transfection via centrifugation at 18,000 x *g* for 15 min. Cell culture supernatants were filtered using a 0.22-μm filter prior to purification.

Filtered Expi cell culture supernatants were buffered with 1/10 volume 10x PBS and loaded onto a HiTrap MabSelect SuRe column (Cytiva) equilibrated in PBS [pH 7.4] using a Cytiva ÄKTA Pure system at a flow rate of 3.5 mL/min. The column was subsequently equilibrated with 5 column volumes PBS [pH 7.4] before elution with 3 column volumes 100 mM glycine [pH 2.8] into 1/10^th^ volume of 1M Tris [pH 8.0]. The column was washed with 0.5 M NaOH with a minimum contact time of 15 min between purifications of different antibodies. Elutions were concentrated using Amicon spin filters (MWCO 10 kDa; Millipore Sigma) and were subsequently loaded onto a GE Superdex S200 increase 10/300 GL column pre-equilibrated in 1X PBS using a Cytiva ÄKTA Pure system. Protein-containing fractions were identified by A280 signal and/or SDS-PAGE, pooled, and stored at 4 °C or at -20 °C in 10% glycerol / 1X PBS until use.

### Env-pseudotyped lentivirus production

Pseudotyped lentivirus bearing various HIV-1 Env proteins were produced by transient transfection of HEK293T cells. For standard TZM-bl assays, the pSG3ΔEnv backbone was used as described previously^29^; for HOS-CD4-CCR5 assays, the pHAGE backbone was used instead as described elsewhere^62^. Virus-containing media were harvested after 2 days, centrifuged, and 0.45-μm filtered. Virus infectivity was titered to ensure that similar levels of infection occurred across viruses and infectivity assays.

### HOS-CD4-CCR5 neutralization assay

HOS-CD4-CCR5 cells (human osteosarcoma cell line stably expressing human CD4, CCR5, and low levels of CXCR4) were plated at a density of 5 × 10^3^ cells per well in white-walled 96-well tissue culture plates (Greiner Bio-One 655098) in growth medium (DMEM, 10% fetal bovine serum (Gemini Bio-Products), 2 mM L-glutamine, 1% penicillin-streptomycin (Sigma-Aldrich), 10 mM HEPES [pH 7.0], 0.22-μm filter-sterilized). The next day, media were removed and replaced with a 100 μL mixture of inhibitor in PBS, HIV-1 pseudotyped lentivirus in growth medium, and growth medium at a final concentration of 2.5 μg/mL DEAE-dextran (Millipore Sigma). After 2 days, 50 μL were aspirated from the plates and 50 μL of BriteLite Luciferase Substrate (Perkin Elmer LLC) were mixed with each well. After 1 min incubation, luminescence was assayed on a Synergy BioHTX plate reader (BioTek). Neutralization IC_50_ values were determined using a three-parameter dose-response curve fit in Prism 9 software (GraphPad).

### TZM-bl neutralization assay

TZM-bl neutralization assays were performed as described previously^29^. Briefly, 5 × 10^3^ TZM-bl cells (HeLa luciferase/β-galactosidase reporter cell line stably expressing human CD4, CCR5, and CXCR4) were plated per well in white-walled 96-well tissue culture plates (Greiner Bio-One 655098) in growth medium (DMEM, 10% fetal bovine serum (Gemini Bio-Products), 2 mM L-glutamine, 1% penicillin-streptomycin (Sigma-Aldrich), 10 mM HEPES [pH 7.0], 0.22-μm filter-sterilized). The next day, media were removed and replaced with a 100 μL mixture of inhibitor in PBS, HIV-1 pseudotyped lentivirus in growth medium, and growth medium at a final concentration of 10 μg/mL DEAE-dextran (Millipore Sigma). After 2 days, 50 μL were aspirated from the plates and 50 μL of BriteLite Luciferase Substrate (Perkin Elmer LLC) were mixed with each well. After 1 min incubation, luminescence was assayed on a Synergy BioHTX plate reader (BioTek). Neutralization IC_50_ values were determined using a three-parameter dose-response curve fit in Prism 9 software (GraphPad).

### Peptide synthesis

CHR peptides used for biolayer interferometry were synthesized by standard Fmoc-based solid-phase peptide synthesis (China Peptides Co. Ltd.). Peptides were modified to contain an N-terminal biotin-PEG6 linker and a C-terminal amide group. Peptides were obtained as a pure lyophilized species and were reconstituted in PBS prior to use.

### Biolayer interferometry (Octet)

Biotinylated peptides (100 nM) were loaded on streptavidin biosensors (Pall ForteBio) to a load threshold of 0.1 nm. Sensors were immediately regenerated in 100 mM glycine [pH 1.5] and neutralized to remove aggregates and non-specific interactions. Ligand-loaded sensors were dipped into known concentrations of 5-Helix for an association step of 60 min. 5-Helix was purified by size-exclusion on the same day as the biolayer interferometry measurements. All reactions were run in PBS with 0.1% BSA, 0.05% Tween 20, and 1 M guanidine hydrochloride. All samples in all experiments were baseline-subtracted to a well that loaded the tip with biotinylated ligand but did not go into sample, as a control for any buffer trends within the samples. The reported K_D_ values were computed using steady state analysis in Octet software (Sartorius) are averaged across experiments repeated on two separate days.

## Abbreviations

bnAbs: broadly neutralizing antibodies
CHR: C-heptad repeat
Env: Envelope
HIV-1: human immunodeficiency virus-1
NHR: N-heptad repeat

## Author Contributions

BNB, TUJB, and PSK conceived experiments and wrote the manuscript; BNB and TUJB performed protein purification/characterization, binding assays, and viral neutralization experiments; NF performed protein purification/characterization. All authors contributed to revising the manuscript.

## Data sharing plan

All data in this manuscript is presented in either main text figures or supplementary material. Unique materials and raw data associated with this manuscript will be readily available upon request.

## Acknowledgments and Funding Information

We thank members of the Kim Lab for critical reading of this manuscript. BNB was supported by the NSF GRFP. TUJB was supported by the Knight-Hennessy Graduate Scholarship and a Canadian Institutes of Health Research Doctoral Foreign Study Award (FRN:170770). This work was supported by the Virginia and D. K. Ludwig Fund for Cancer Research, the Chan Zuckerberg Biohub, and an NIH Director’s Pioneer Award (DP1DA043893) to PSK. Figure 1 and Supplementary Figure 1 were created using Biorender.

